# CellTool: an open source software combining bio-image analysis and mathematical modeling

**DOI:** 10.1101/454256

**Authors:** Georgi Danovski, Teodora Dyankova, Stoyno Stoynov

## Abstract

**Summary:** We present CellTool, a stand-alone open source software with a Graphical User Interface for image analysis, optimized for measurement of time-lapse microscopy images. It combines data management, image processing, mathematical modeling and graphical presentation of data in a single package. Multiple image filters, segmentation and particle tracking algorithms, combined with direct visualization of the obtained results make CellTool an ideal application for rapid execution of complex tasks. In addition, the software allows for the fitting of the obtained results to predefined or custom mathematical models. Importantly, CellTool provides a platform for easy implementation of custom image analysis packages written on a variety of programing languages.

**Availability and Implementation:** CellTool is a free software available for MS Windows OS under the terms of the GNU General Public License. Executables and source files, supplementary information and sample data sets are freely available for download at URL: https://dnarepair.bas.bg/software/CellTool/

**Supplementary information:** Supplementary data are available at URL: https://dnarepair.bas.bg/software/CellTool/Program/CellTool_UserGuide.pdf

## Introduction

Present-day microscopy and protein labeling techniques allow for fast recording of rapid processes in living cells (Wu, et al., 2011). The acquired large datasets are currently analyzed by using multiple software packages for image processing, data management and mathematical modeling, which is difficult and time consuming (Swedlow, et al., 2003). Moreover, accurate measurement of the objects in living systems requires testing of different segmentation and tracking algorithms with fine tuning of their settings, considerably slowing down the process of image analysis. Smooth image analysis workflow requires a software that implements constant feedback between image data processing and graphical data presentation.

## Results and discussion

### Program overview

Herein, we present CellTool, a stand-alone open source software highly optimized for parallel computing that combines tools for image processing, data management, graphical data presentation and mathematical modeling in a single package. It has implemented algorithms for fast, automatic segmentation of large time-lapse image stacks and a newly developed algorithm for particle detection. There are various image filters and binary operations that can be applied to improve the quality of the segmentation. All of these settings can be stored as protocols and automatically applied. Different types of regions of interest (ROIs) can be used in order to measure particular areas of the image. We developed a new algorithm that is capable of tracking and measuring the objects of interest over time. The acquired data is directly visualized on a chart and can be viewed or modified by applying custom functions, which saves time and eliminates the step of post-processing. The summarized results from the analysis of the image series can be normalized, filtered and graphically presented with the “Results Extractor” plugin in CellTool. Importantly, the plugin can be used for curve fitting of the raw data to predefined or custom mathematical models. Such functionality allows for scientists without strong mathematical background to perform mathematical modeling of their results.

The user-friendly graphical interface of CellTool enables constant feedback between the different tasks to ensure fast and accurate image analysis (Fig 1A). A generalized workflow in CellTool includes the following steps (for a full description, manual of use and definitions, see Supplementary Material).

1. The user can add a work directory to the “Data Sources” panel. This integrated file manager is created for fast and easy access to the data. From there an image or a group of images can be opened in new tabs in the central “work” panel.
2. The image could be edited via different pre-processing steps including making substacks, projections, splitting, merging channels or cropping.
3. By clicking on the “Processed image”, different settings are enabled. Filters and binary operations can be applied and the image can be segmented using global methods of segmentation and particle detection.
4. Two types of ROIs, “Tracking” and “Static”, can be used to measure the objects. Different settings for the ROIs can be applied, including a variety of shapes and a multilayer option.
5. The obtained results from the ROIs can be immediately viewed on the “Results chart” and can be exported as a tab-delimited text file. Here a predefined formula can be used to eliminate some of the post-processing steps.
6. Filtering, normalization and visualization of the data from an image series can be performed in the “Results Extractor” plugin.
7. Batch curve fitting of the acquired data to custom or predefined mathematical models can be also done in the “Results Extractor”.

**Figure 1.**
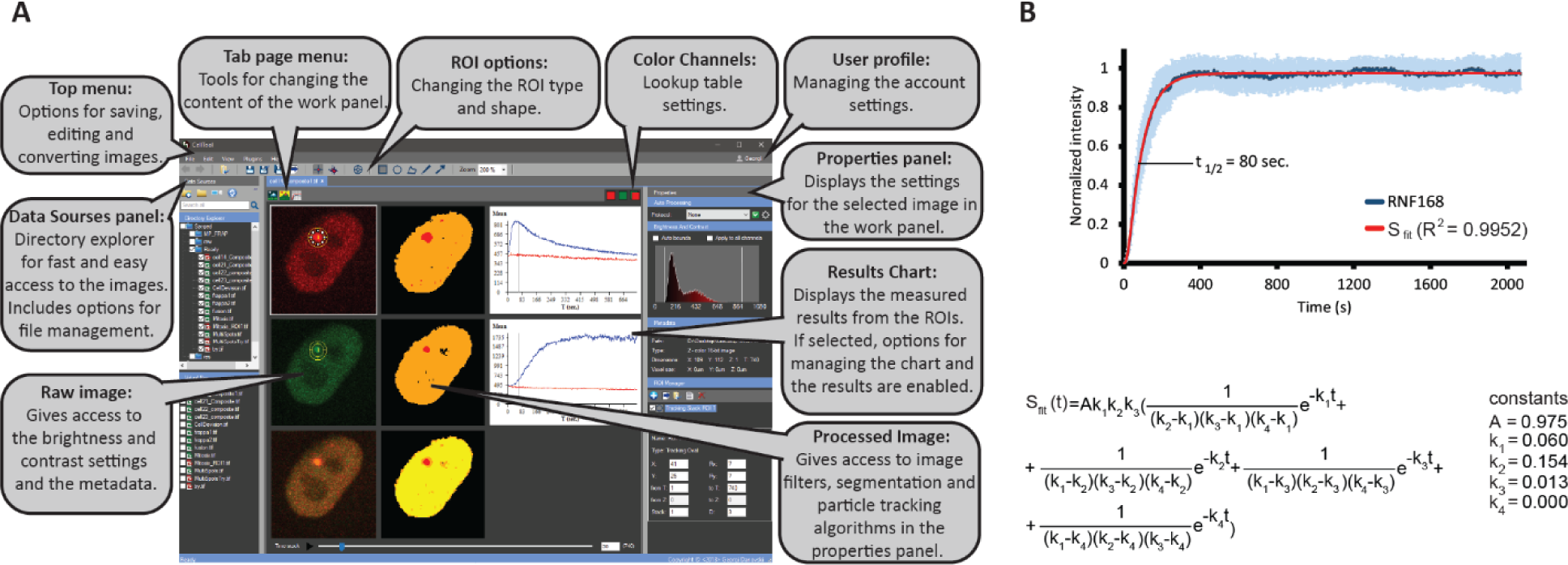
**(A)** CellTool’s main Graphical User Interface. **(B)** Kinetic of recruitment of RNF168 to DNA damage sites and curve fitting results

Our software is able to open most of the available formats for microscopy images (Linkert, et al., 2010). CellTool supports 8-bit grayscale images and 16-bit grayscale images (RGB images are split into 3 separate grayscale images). Additionally, the file metadata can be reviewed.

Furthermore, there is a plugin platform which allows for new image analysis algorithms to be added to CellTool. The plugins must be built as a Class Library and written in one of the following Microsoft ․NET Framework languages: C#, Visual Basic or F# (see Supplementary Material).

CellTool has a password protected multi-user interface under the control of an administrator. The settings for each user are stored and will be automatically uploaded upon login. This feature makes CellTool suitable for microscopy imaging facilities.

In summary, CellTool is a novel open source software with highly intuitive graphical interface capable of semi-automatic image processing, mathematical modeling and simultaneous data visualization, which greatly increases the efficiency of the image analysis. CellTool is perfect for rapid and robust live cell image data analysis providing an all-in-one solution to numerous demanding tasks.

### Test case

CellTool was used for the measurement and modeling of the kinetic of recruitment of EGFP-tagged RNF168 at complex DNA damage sites (Aleksandrov, et al., 2018) (Fig 1B). The images are obtained with a spinning disk confocal microscope with a UV laser module for microirradiation. Detailed image analysis workflow is available in the Supplementary Material. Our analysis reveals that the halftime of recruitment of RNF168 at the sites of complex DNA damages is 80 sec. The halftimes of recruitment and removal provide a fast and simple way to quantify the kinetics of DNA repair proteins. However, the halftimes of recruitment cannot distinguish a single-step repair reaction from a more complex, multi-step engagement of the protein in the repair process. Therefore, we fit the total intensity data for RNF168 to a Consecutive Reactions Chain (CRC) model (Aleksandrov, et al., 2018) that describes protein recruitment to damage sites as a series of consecutive reactions followed by a period during which the proteins reside on the lesions and a final reaction responsible for the removal of the proteins from damaged regions (Fig 1B). The modeling shows that RNF168 is recruited to the sites of DNA damage by at least 3 consecutive reactions and reveals their kinetic parameters (Aleksandrov, et al., 2018). The results demonstrate the ability of our software not only to rapidly process and measure image data but also to reveal the underlying mechanisms of the studied cellular processes by mathematical modeling.

## Acknowledgements

*Funding*: We gratefully acknowledge the funding from the National Science Fund of Bulgaria (ДН11/17 and ДН11/8) and ICGEB (CRP/BGR16-03).

*Conflict of Interest*: none declared.

